# Back from the dead: the atypical kinase activity of a pseudokinase regulator of cation fluxes during inducible immunity

**DOI:** 10.1101/2022.05.10.491368

**Authors:** Elizabeth K. Brauer, Nagib Ahsan, George V. Popescu, Jay J. Thelen, Sorina C. Popescu

## Abstract

While most pseudokinases lack phosphotransfer activity due to altered catalytic residues within their kinase domain, a subset of pseudokinases maintain kinase activity through atypical mechanisms. The Arabidopsis ILK1 is a pseudokinase from the Raf-like MAP3K family and is the only known plant pseudokinase with confirmed kinase activity. ILK1 activity promotes microbial pattern-induced root growth inhibition through its stabilization of the HAK5 potassium transporter with the calmodulin-like protein CML9. ILK1 also has a kinase-independent function in salt stress suggesting that it interacts with additional proteins. We determined that members of the ILK subfamily are the sole pseudokinases within the Raf-like MAP3K family and identified 179 novel putative ILK1 protein interactors. We also identified 70 novel peptide targets for ILK1, the majority of which were phosphorylated in the presence of Mn^2+^ instead of Mg^2+^ in line with modifications in ILK1’s DFG cofactor binding domain. Overall, the ILK1-targeted or interacting proteins included diverse protein types including transporters (HAK5, STP1), protein kinases (MEKK1, MEKK3), and a cytokinin receptor (AHK2). The expression of 31 genes encoding putative ILK1-interacting or phosphorylated proteins, including AHK2, were altered in the root and shoot in response to molecular patterns suggesting a role for these genes in immunity. We describe a potential role for ILK1 interactors in the context of cation-dependent immune signaling responses, highlighting the importance of K^+^ in MAMP responses. This work further supports the notion that ILK1 is an atypical kinase with an unusual cofactor dependence that may interact with multiple proteins in the cell.

## Introduction

Protein kinases form the basis of the phosphorylation cascades that transmit cellular signals by facilitating phosphotransfer from ATP to serine, threonine, tyrosine, and histidine residues in a substrate protein or peptide (Kwon et al., 2019). The highly conserved eukaryotic kinase domain contains three catalytic residues essential for binding and positioning ATP in conjunction with metal ion cofactors such as Mn^2+^ and Mg^2+^ to facilitate phosphotransfer (Adams, 2001). Approximately 5% of all plant genes encode proteins containing a kinase domain, and one-third of plant proteins are phosphorylated (Kwon et al., 2019). Based on bioinformatic analyses, eukaryotic kinases have been divided into canonical (catalytically active) kinases and pseudokinases, which are usually catalytically inactive due to a non-canonical catalytic residue (Kwon et al., 2019). The catalytic residues include aspartate in the DFG subdomain that binds divalent cations to co-ordinate β and γ phosphates of ATP, the lysine in the VAIK subdomain that anchors and orients ATP for substrate transfer and the aspartate in the HRD domain of the catalytic loop that is a base acceptor to achieve proton transfer (Adams, 2001). Over 17% of predicted protein kinases encoded in the Arabidopsis genome are classified as peudokinases and recent research on plant pseudokinases confirm that they perform distinct biological functions in plant defense against microbial pathogens, which rely on protein-protein interactions, as opposed to canonical catalysis (Kwon et al., 2019). For example, several kinase catalytic subdomains of the ZED1 pseudokinase influences its binding to PBL proteins and the ZAR1 nucleotide-binding leucine-rich repeat (NLR) protein during activation of effector-triggered immunity (ETI) (Bastedo et al., 2019). The BIR2 pseudokinase regulates microbial-associated molecular pattern (MAMP)-induced signaling by binding the BAK1 co-receptor to prevent its untimely interaction with the FLS2 receptor which is involved in perception of the flagellin-derived flg22 (Halter et al., 2014). The GHR1 receptor kinase is involved in inducing stomatal closure following MAMP treatment through its interactions with the calcium-responsive CDPK3 (Sierla et al., 2018; Hua et al., 2012).

While a majority of pseudokinases lack phospho-transfer activity, a small subset maintain atypical kinase activity through substitutions with amino acids elsewhere in the catalytic site to facilitate phosphotransfer (Kwon et al., 2019). For example, some pseudokinases do not require a cofactor to initiate phosphotransfer indicating the divergence of their phosphotransfer mechanisms from typical protein kinases (Mukherjee et al., 2010; Kwon et al., 2019; Lewis et al., 2014). The integrin-liked kinase 1 (ILK1) is the only plant pseudokinase which is an active kinase *in vitro* and *in vivo* with demonstrated auto- and substrate-phosphorylation activity (Brauer et al., 2016, Nemoto et al., 2011). ILK1 belongs to the ILK family of Raf-like MAP3Ks found in both metazoans and plants though the nature of kinase catalytic residue degeneration is distinct between kingdoms (Popescu et al., 2017, Ichimura et al., 2002). In animals, the ILKs lack the catalytic aspartate in the HRD domain and are likely inactive, though some evidence suggests that they maintain kinase activity *in vitro* (Hannigan et al., 2011; Dagnino, 2011). Plants lack integrin homologs and typically encode ILKs with both canonical kinase domains and pseudokinases containing a GFG motif in place of the metal ion cofactor binding DFG motif (Popescu et al., 2017). In Arabidopsis, *ILK1, ILK2* and *ILK3* encode pseudokinases while *ILK4, ILK5* and *ILK6* encode kinases. Recent work indicates that ILK5 is phosphorylated by the extracellular ATP receptor P2K1 and phosphorylates MKK5 to activate MPK3 and MPK6 (Kim et al., 2022a). The ILK5-MKK5-MPK3/6 cascade induces stomatal closure and resistance to *Pseudomonas syringae* pv. *tomato* (DC3000) (Kim et al., 2022a). ILK6 was identified as a putative interactor of the VH1 BRI1 family receptor-like kinase and *ilk6* mutants display altered vein pattern formation in cotyledons (Ceserani et al., 2009). Both *ILK5* and *ILK6* promote stomatal closure in response to blue light (Hayashi et al., 2017). Interestingly, two plant pseudokinase ILKs from Arabidopsis (ILK1) and Medicago (MsILK1) are able to phosphorylate substrates *in vitro* and have an unusual preference for the Mn^2+^ cofactor compared to Mg^2+^ (Brauer et al., 2016; Chinchilla et al., 2008). Kinase activity of the Arabidopsis ILK1 is essential for root growth inhibition in response to the flg22, pep1, and elf18 molecular patterns and resistance to an effectorless DC3000 pathogen (Brauer et al., 2016; Popescu et al., 2017). The role of ILK1 in defense seems to be partly dependent on its maintenance of HAK5 potassium transporter stability through interactions in conjunction with the calmodulin-like protein CML9 (Brauer et al., 2016). While mutation of the aspartate residue in the HRD catalytic loop to asparagine eliminates kinase activity in most protein kinases, this mutation increased ILK1 activity further demonstrating the uniqueness of its enzymatic mechanism (Strong et al., 2011; Brauer et al., 2016). Ectopic expression of the Medicago ILK1 containing the same HRD->HRN mutation in Arabidopsis reduced lateral root formation relative to the native Medicago ILK1, further indicating a role for the kinase activity in root development (Chinchilla et al., 2008). Kinase-independent functions for ILK1 include the promotion of germination during salt and osmotic stresses suggesting that much like the metazoan ILKs that scaffold multiple proteins to promote cell adhesion, ILK1 may also interact with more than one protein *in vivo* (Brauer et al., 2016, Chastney et al., 2021). Understanding the capacity of the plant ILKs to interact and phosphorylate targets would expand our comprehension of the active pseudokinases and facilitate future work on modeling alternative active sites and prediction of protein targets. Towards this goal, we used *in vitro* approaches to identify ILK1 interacting proteins and phosphorylated peptides and determine how the Mn^2+^ or Mg^2+^ cofactors influenced the activity of the protein *in vitro*.

## Results

### Putative ILK1 phosphorylation targets

Our previous survey of plant ILKs indicates the DFG->GFG substitution in the catalytic site occurred prior to the divergence between the gymnosperms and the angiosperms (Popescu et al., 2017). We surveyed variation within the three essential catalytic residues to determine if similar substitutions exist across the Raf-like MAP3K family. The ILKs were the only pseudokinases within the Raf-like MAP3Ks suggesting that the GFG substitution of ILKs developed independently during evolution (Supplementary Table 1). The ILKs are also non-RD kinases (HRD-> HCD) and are overrepresented alongside the B4 subfamily as non-RD kinases within Raf-like MAP3K family (Supplementary Table 1). Non-RD kinases do not require activation loop phosphorylation to become active and can instead be activated through other conformational changes and autophosphorylation sites in ILK1 are indeed outside of activation loop (Nolan et al., 2004, Brauer et al., 2016).

To further characterize the unusual kinase activity of ILK1, we identified phosphorylation targets of ILK1 using the kinase client (KiC) assay. This approach involves incubation of a purified protein kinase with a synthetic peptide library comprised of approximately 2,100 peptides, representing over 4,000 *in vivo*, experimentally-mapped phosphorylation sites, and detection of phosphorylated sites by mass spectrometry (Ahsan et al., 2013, Huang and Thelen, 2012, Huang et al., 2010). We identified 32 and 18 phosphorylated peptides using either Mn^2+^ or Mg^2+^ as cofactors, respectively (Figure 1, Supplementary Tables 2, 3, Supplemental Data 1). Four peptides were phosphorylated in both conditions, including one derived from a pectin methylesterase involved in cell wall-mediated defense (PME39, AT4G02300), the BYPASS1 protein which is required for root responses to *Rhizobium* infection, an ankyrin repeat protein (AT3G04470) and an RNA helicase family protein (AT1G08050) that is oxidized following flg22 or salicylic acid treatments (Bethke et al., 2009; Liu et al., 2015; Arthikala et al., 2018). The majority of the ILK1 peptide targets were localized to the nucleus or plasma membrane, consistent with previous observations that ILK1 can be found in the plasma membrane, the endoplasmic reticulum, and the nucleus (Brauer et al., 2016)(Supplementary Figure 1A, Supplemental Data 1). Applying the KiC approach using the hyperactive ILK1^D319N^ isoform revealed 26 and 16 phosphorylated peptides with Mn^2+^ or Mg^2+^ respectively, 15 of which were also identified as phosphorylated by ILK1. Altogether, 70 peptide targets were phosphorylated by ILK1 and ILK1^D319N^ including transporters, protein kinases, enzymes involved in metabolite biosynthesis, and scaffold proteins (Supplementary Figure 1B). Notable potential targets included the HAK5 transporter which was confirmed as an ILK1-interacting protein previously, MEKK1, which promotes basal defense downstream of PRRs and MEKK7, which interacts with the FLS2 PRR to suppress flg22-induced immune responses (Brauer et al., 2016, Sun and Zhang, 2022; Mithoe et al., 2016).

**Figure 1:**
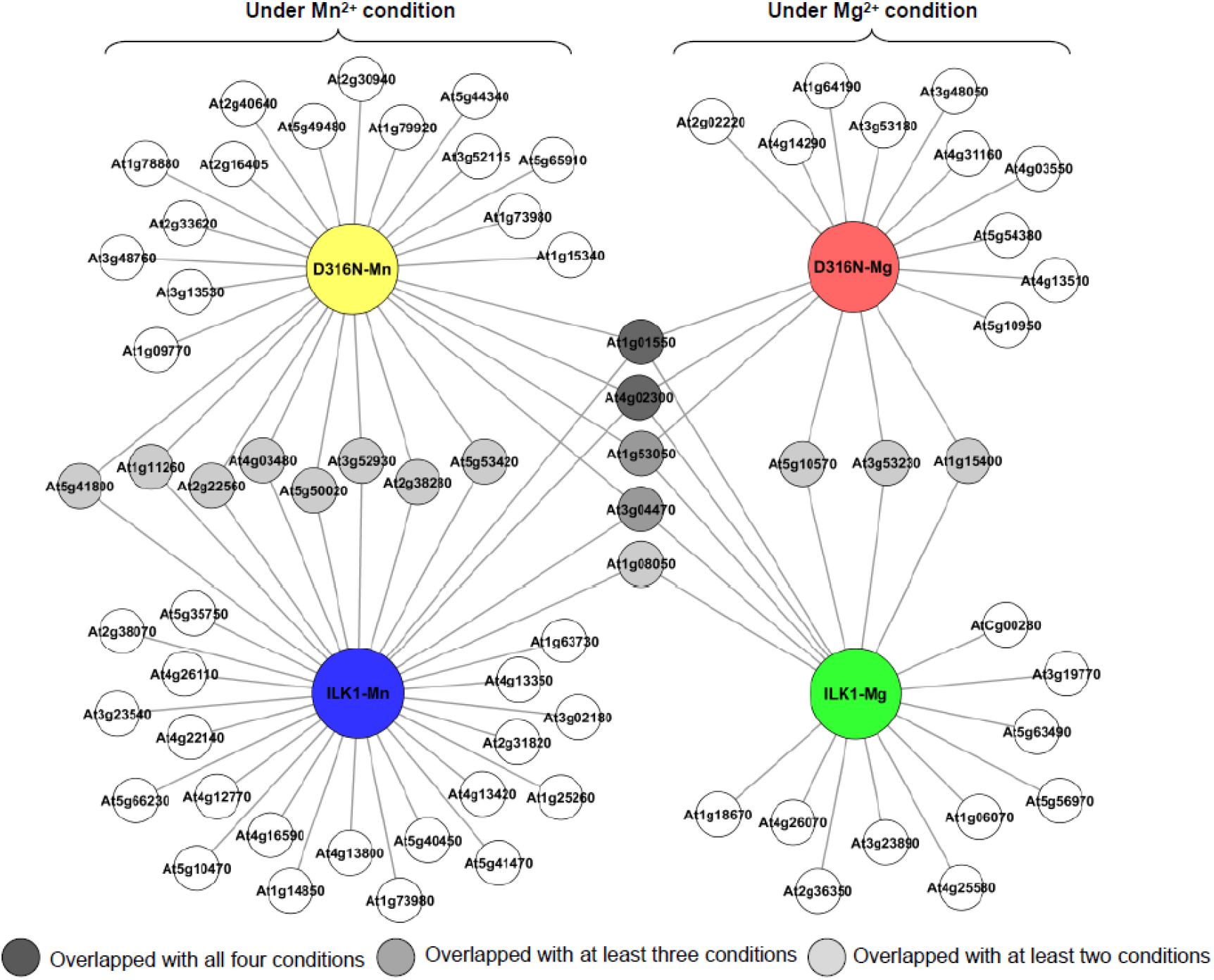
Topological relationship of the potential phosphorylation targets for ILK1 and the ILK^D316N^ hyperactive mutant identified by screening the phosphorylation status of a synthetic peptide library after incubation with purified ILK1 protein. Cofactor conditions included the use of either 5 mM MgCl_2_ or MnCl_2_. The cartograph was assembled by Cytoscape 2.8.3 (http://www.cytoscape.org).

In general, both ILK1 and ILK1^D319N^ kinases phosphorylated primarily serine and threonine residues, but tyrosine phosphorylation was also recorded on a small subset of peptides, including one derived from the HAK5 transporter (Supplementary Table 2,3). A higher number of targets was observed with Mn^2+^ as a cofactor compared to Mg^2+^ which is interesting considering that Mg^2+^ is preferred by most serine/threonine protein kinases while tyrosine kinases seem to prefer Mn^2+^ as a cofactor (Swarup et al., 1984; Bossemeyer et al., 1993). Thus, ILK1 appears to be an exception to these trends, being primarily a serine/threonine kinase with a preference for the Mn^2+^ cofactor.

### Putative ILK1 protein interactors

To understand the protein-protein interaction specificity of ILK1, we surveyed half (15,000) of the predicted Arabidopsis proteome using functional protein microarrays to identify 179 putative interactors (Supplemental Data 2). Most of the interactors localize to the nucleus or the cytosol, and are involved in metabolite conversion, protein modification, or transcriptional regulation (Supplementary Figure 1, Supplemental Data 2). Interactors within the MAPK family included the MEKK3 that promotes MTI and signal transduction downstream of PRR activation (Sun and Zhang, 2022). We also identified MEKK19, and MPK19 as potential interactors, though the function of these kinases is unknown. Several transporters were also identified as potential ILK1 interactors including the KUP5 potassium transporter which is closely related to HAK5, and is expressed across diverse root cell types (Lhamo and Luan, 2021). This indicates that ILK1 can interact with a wide range of protein types *in vitro*, supporting the notion that additional ILK1-protein interactions beyond the CML9 and HAK5 proteins may exist in plant cells.

### Trends across putative ILK1 protein interactors and substrates

Overall, we identified 249 peptides or proteins which interacted with purified ILK1 *in vitro* and are not significantly enriched for specific biological or molecular functions. Interestingly, we did not identify the same ILK1 targets in both the KiC and functional protein microarray, though these approaches shared 709 peptides or proteins derived from the same cognate protein (Ahsan et al., 2013; Popescu et al., 2007). Within the shared protein list, 28 ILK1-phosphorylated peptides had their cognate proteins spotted on the functional microarray but were not identified as ILK1-interacting proteins on the microarray. These included the HAK5 transporter that interacts with ILK1 *in vivo*, the MEKK1 immune signaling kinase, the STP1 sugar transporter, and the THE1 receptor kinase that detects cell wall modifications to promote resistance against necrotrophic pathogens (Brauer et al., 2016, Qu et al., 2017). This would suggest that protein microarrays may generate false negatives likely due to the influence of the interaction surface on the protein folding and protein-protein interactions. While further work is needed to confirm these interactions, combining proteomic approaches enabled us to assemble a calcium-responsive complex between CML9, ILK1 and HAK5 in addition to confirmed substrates for the P2K1 receptor kinase (Popescu et al., 2007, Brauer et al., 2016, Chen et al., 2017, Chen et al., 2021). Further, the genes transcribing 31 of the putative ILK1 interactors were transcriptionally responsive to MAMP treatments in roots or shoots suggesting a role in immunity (Figure 2, Supplementary Table 4). These included three transporters, the cytokinin receptor AHK2 and the pectinesterase inhibitor PMEI4, both of which influence root growth (Kubiasová et al., 2020; Hocq et al., 2017). Thus, ILK1’s influence on MAMP-induced root growth inhibition may be linked to its effect on nutrient transport, cell wall modifications or cytokinin signaling and further functional work is needed to explore these possibilities.

**Figure 2:**
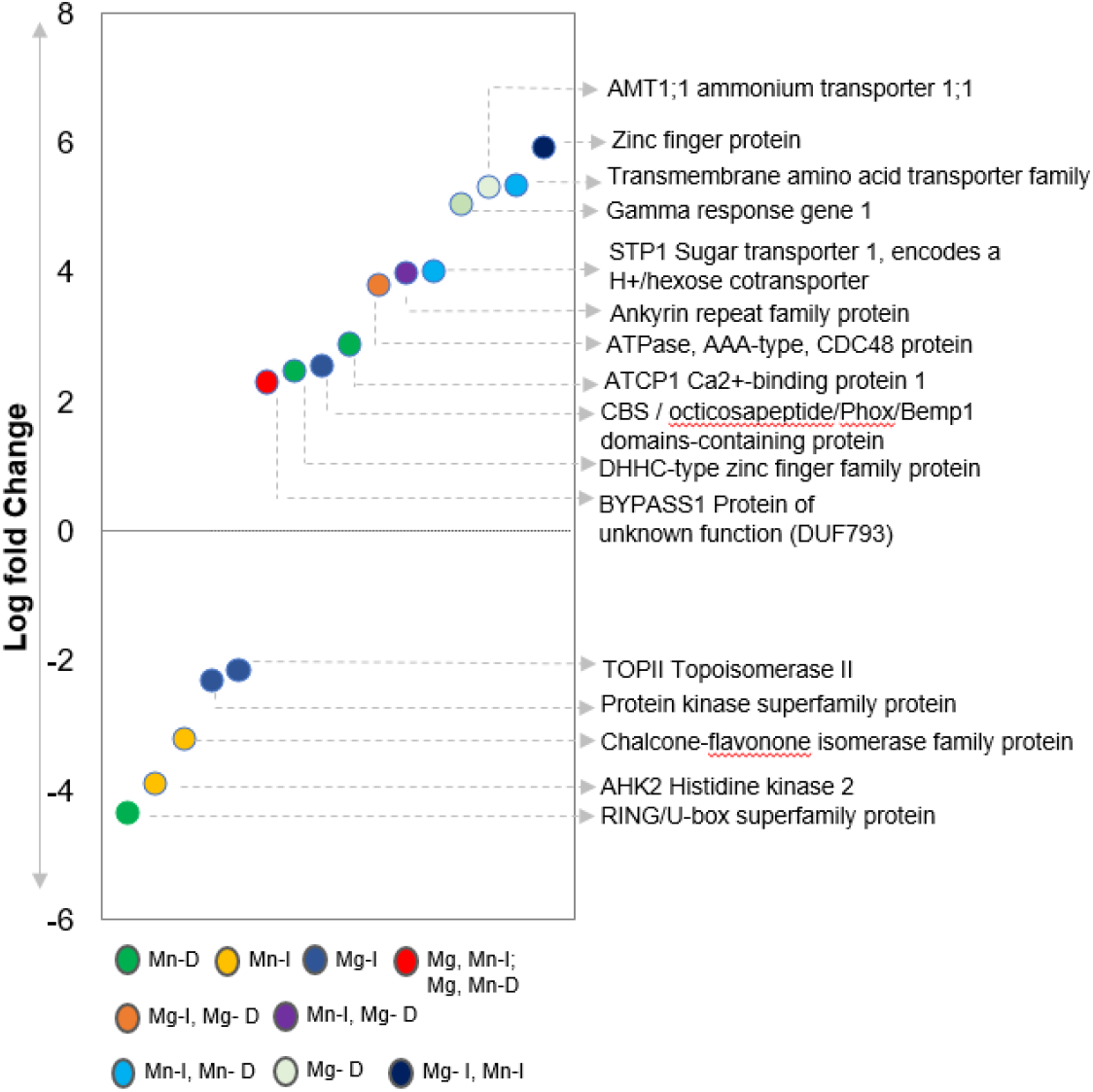
Differential expression of ILK1 targets in Arabidopsis leaves 3 hours after 1uM flg22 treatment relative to the 0 hour time point (from Dimlioglu et al). All of the targets were identified by KiC in the presence of ILK1-V5 (I) of ILK1^D319N^-V5 (D) purified proteins with MnCl_2_ (Mn) or MgCl_2_ (Mg) as cofactors.

## Discussion

Here, we describe the putative protein-protein interactions and phosphorylation activity of the atypical protein kinase ILK1. Members of the ILK family were the only pseudokinases identified within the Raf-like MAP3Ks and the ILK1 is one of the only pseudokinases in plants with demonstrated phosphotransfer activity demonstrated *in vitro* and *in vivo* (Brauer et al., 2016). In line with its unusual metal ion cofactor-binding domain, we demonstrate that ILK1 phosphorylates a broader range of peptides in the presence of Mn^2+^ compared to Mg^2+^ but that it primarily phosphorylates serine and threonine residues. Our survey of ILK1 protein interactors revealed interactions with diverse proteins, including transporters, protein kinases, cell wall modifying enzymes, and hormone receptors. Together, this indicates that ILK1 is an active kinase with a potentially broad range of interactions with diverse substrates. While its kinase-dependent role in MAMP-induced growth and defense responses seems to align with its regulation of the HAK5 potassium transporter, we cannot exclude the possibility that additional phosphorylation targets of ILK1 may exist in the cell. Indeed, it is increasingly apparent that protein kinases interact with a wide range of proteins *in vitro* and *in vivo*, suggesting that the functional specificity of kinases is related to their localization, protein complexing, and posttranslational modifications as opposed to interaction specificities (Popescu et al., 2009; Brauer et al., 2018).

Root growth is influenced by both potassium and MAMPs through ILK1 and HAK5 indicating shared components of potassium and immune signaling pathways (Brauer et al., 2016, Qi et al., 2008). Blocking K^+^ channels abolishes flg22-induced root growth inhibition which is counterintuitive considering that reducing K^+^ uptake should reduce root growth instead of promoting it (Brauer et al., 2016). The importance of potassium in defense is further underlined by the fact that the rice blast pathogen *Magnaporthe oryzae* targets the AKT1 potassium channel using the AvrPiz-t effector to promote virulence during infection (Shi et al., 2018). Bulk Ca^2+^, K^+^ loss and P and S gain in Arabidopsis seedlings is induced within minutes of flg22 exposure and this shift in nutrient homeostasis is reversed within 3 hours (Brauer et al., 2016). On the pathogen side, *P. syringae* DC3000 sulfur starvation responses are differently induced when plants are pre-treated with flg22 compared to no pretreatment indicating that PTI influenced nutritional stresses imposed on the pathogen (Lovelace et al., 2018).

At the cellular level, MAMP-induced cation (Ca^2+^, K^+^) transport is transient and contributes to the rate but not the extent of membrane depolarization which is primarily mediated by anions (Brauer et al., 2016; Jeworutzki et al., 2010; Jabs et al., 1997). Cation fluxes through La^3+^-sensitive channels are essential to trigger MAMP-triggered plasma membrane depolarization and defense compound production (Jeworutzki et al., 2010, Jabs et al., 1997). While these effects are attributed to Ca^2+^-depending signaling, La^3+^ blocks both Ca^2+^ and K^+^ channels in plants and thus a more nuanced understanding of the relationship between cation fluxes and immune signaling is warranted (Terry et al., 1992). For example, K^+^ levels influence the sensitivity of the plasma membrane to flg22, potentially through its influence on membrane polarity (Chi et al., 2021). Mn^2+^ homeostasis also changes in response to flg22 which could impact cofactor availability for ILK1 as well as tyrosine phosphorylation of pattern recognition receptors such as the lipid MAMP-binding LORE receptor and the elf18-binding EFR receptor (Macho et al., 2015, Luo et al., 2020). In mammals, cation leakage from pore-forming toxins or atypical cation pore formation is detected by the cell to induce MAPK signaling (Olabisi et al., 2016, Kloft et al., 2009). In plants, recent work indicates that the NB-LRRs form cation-conducting channels in the plasma membrane upon avirulence factor recognition to activate ETI (Kim et al., 2022b). These resistosome channels can conduct calcium, but may also conduct other cations (Kim et al., 2022b). The signaling mechanisms connecting cation efflux with downstream activation of ETI signaling remains unclear but could involve similar mechanisms as PTI. For example, cation-induced changes in plasma membrane polarity influences the voltage-regulated vacuolar two-pore channels and potassium channels including GORK and SKOR (Kurusu et al., 2005; Wu et al., 2018). Membrane polarity also influences membrane fluidity and microdomain formation in which receptors for PTI and ETI activation can be found (Malinsky et al., 2016). Alternatively, cations may be sensed directly as has been recently demonstrated during salt stress, where GIPC sphingolipids bind Na^+^ ions to activate Ca^2+^ influx channels and downstream signaling pathways regulating root growth (Jiang et al., 2019). Our understanding of the influence of induced cation fluxes on inducible immunity remains unclear, but the recent breakthroughs in our understanding of resistosomes will no doubt advance our progress in this area.

This work improves our understanding of the atypical protein kinases and reinforces the notion that functionally relevant alternative protein kinase mechanisms exist. Further characterization of the ILK family and their role as either scaffolding proteins or protein kinases will help to illuminate the function of these pseudokinases in plant defense and stress responses.

## Methods

### Protein purification and protein microarray

The ILK1-V5 and ILK1D319N-V5 proteins were purified as described previously and protein purity was validated by mass spectrometry indicating a lack of background kinases within the samples (Brauer et al., 2014, 2016). Both Mn^2+^ and Mg^2+^ (5 mM) were included during ILK1 incubation on the arrays as cofactors and the experiment was conducted in triplicate as described in Brauer, 2016. Briefly, protein microarrays (AtPMA-5000 and AtPMA-10000) were blocked in TBS-T (20 mM Tris, 137 mM NaCl, 0.1 % (v/v) Tween-20) with 1 % (w/v) BSA for 1 h, drained, and kept in a humid chamber. 150 µl of purified ILK1 or cleavage buffer (both mixed with 1 % (w/v) BSA and 5 mM MgCl2 and MnCl2) were applied to the slides for 1.5 h. Slides were washed in washing buffer (PBS, 5 mM MgCl2 and MnCl2, 0.05 % (v/v) Triton X-100, 5 % (v/v) glycerol, 1 % (w/v) BSA), incubated with 500 µl Cy5-conjugated anti-V5 antibody (1:8,000, Agrisera) for 30 min, washed and incubated with 500 µl α-rabbit-DyLight649 (1:10,000, Jackson Immuno Research) for 30 min and washed with the same buffer once again. Slides were dipped in water, spun-dried, and scanned in a Scanarray® express scanner (Perkin Elmer); images were processed using GenePix® Pro 7 Acquisition and Analysis Software (Molecular Devices) as described in (Popescu et al. 2009). We performed three experiments using ILK1 as a probe and two experiments without probe as a control. Protein interactors with significantly higher V5 signal above background levels across all three experiments were designated as putative ILK1 interactors.

### Protein microarray bioinformatics

To analyze candidates with stronger binding signals than the control we implemented in Matlab a cross-array normalization method followed by a statistical testing method. All microarrays (probe and control) datasets were normalized using a linear regression model as follows: 1) background median intensity was subtracted from signal median intensity; 2) linear regression coefficients for each probe and control pairs were calculated: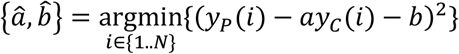, where *N* is the number of proteins printed on the microarray; 3) “between arrays” normalization was performed using a linear transformation of probe datasets (using the estimated regression coefficients) and scaling of control datasets: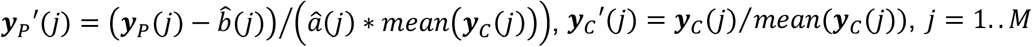, the number of (probe, control) pair datasets. To select probe binding candidates we used a one-side t-test: = 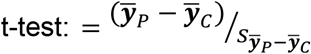, where 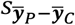 is the standard error of the difference between the means; using pooled variance and 5% significance level (due to the small sample set). We controlled type II errors by applying an FDR method described (Storey, 2002).

### KiC

The same ILK1 purified proteins that were used for the protein microarray were also used for the KiC as described previously (Ahsan et al., 2013; Brauer et al., 2016). Peptide spectral matches were evaluated primarily using the XCorr scoring function of SEQUEST, employing a 1% false discovery rate for a randomized database of the peptide library. Phosphorylation-site localization was performed using phosphoRS (pRS) (Proteome Discoverer, v1.0.3, Thermo Fisher). An empty vector was used as a control against the 2.1k peptide library, peptides that overlapped with the empty vector control were excluded as potential clients, as these likely represented “background” kinase activities as described in Brauer 2016.

### Annotation and enrichment analysis

To determine functional enrichment for ILK1 interactors or ILK1 phosphorylation targets within a dataset, the PANTHER GO Term Enrichment tool (http://pantherdb.org/) was used with a cut-off value of p<0.05. Enrichment was measured against both the entire Arabidopsis genome as the background or using the genes specifically present in the KIC library or protein microarray.

## Supporting information

KIA interactors

FPM interactors

