## Supplemental Tables and Figures for "Back from the dead: the atypical kinase activity of a pseudokinase regulator of cation fluxes during inducible immunity"

**Supplementary Files:**

**Supplementary Table 1**: Analysis of catalytic residues in the kinase domains of the Arabidopsis Raf-like MAP3Ks as defined in Ichimura et al., 2002.

| Raf-like MAP3K subgroup | N^1^ | VAI**K**^2^ | H**R**D^2^ | HR**D**^2^ | **D**FG^2^ |
| --- | --- | --- | --- | --- | --- |
| B1 | 3 | 0 | 0 | 0 | 0 |
| B2 | 6 | 0 | 0 | 0 | 0 |
| B3 | 6 | 0 | 0 | 0 | 0 |
| B4 | 7 | 0 | 7 | 0 | 0 |
| C1  (ILK family) | 6 | 0 | 4 | 1 (ILK2) | 3 (ILK1,2,3) |
| C2 | 3 | 0 | 0 | 0 | 0 |
| C3 | 4 | 0 | 0 | 0 | 0 |
| C4 | 3 | 0 | 0 | 0 | 0 |
| C5 | 3 | 0 | 1 | 0 | 0 |
| C6 | 2 | 0 | 0 | 0 | 0 |
| C7 | 5 | 0 | 0 | 0 | 0 |
| Total | 48 | 0 | 12 | 1 | 3 |

^1^Number of genes within the subgroup.

^2^Number of members lacking the catalytic residue underlined and indicated in bold within the indicated conserved subdomain. Note the arginine within the HRD subdomain is not essential for phosphotransfer but is of interest due to its association with kinase activation mechanisms.

**Supplementary Table 2:** Putative kinase clients identified by KiC assay using a synthetic peptide library consisting 2.1K peptides as substrates for *Arabidopsis thaliana* ILK1 (At2G43850) kinase and its D319N mutant in presence of Mg (5 mM).

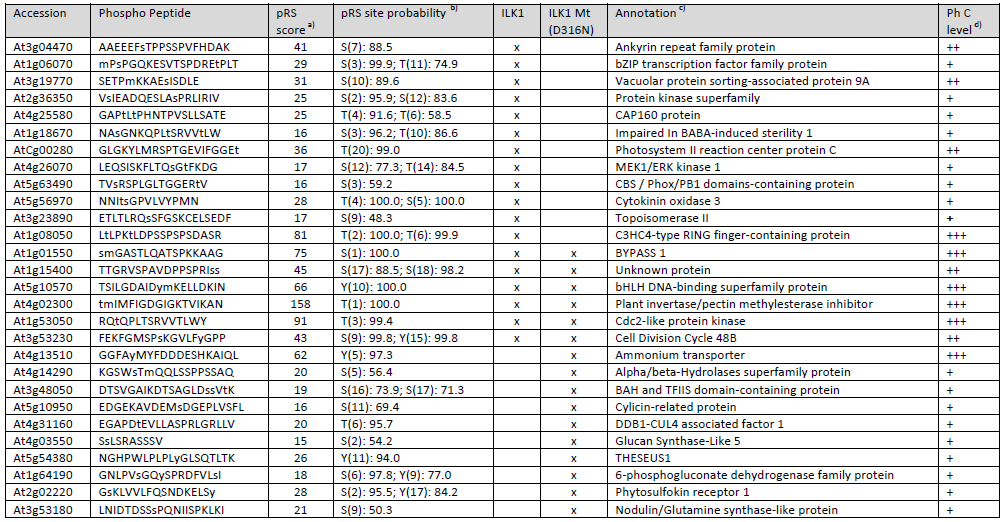

a) pRS score: This peptide score is based on the cumulative binomial probability that the observed match is a random event. The value of the pRS score strongly depends on the data scored, but usually scores above 50 give good evidence for a good PSM.

b) pRS site Probabilities: For each phosphorylation site this is an estimation of the probability (0-100%) for the respective site being truly phosphorylated. pRS Site Probabilities above 75% are good evidence that the respective site is truly phosphorylated

c) Protein annotation from TAIR database

d) Ph C level: Confidence level of the phosphopeptides

+++ (High): pRS score >50; pRS site probability >75%;

++ (Moderate): pRS score 30-49; pRS site probability 40-74%;

+ (Low): pRS score 15-30; pRS site probability >40%;

**Supplementary Table 3:** Putative kinase clients identified by KiC assay using a synthetic peptide library consisting 2.1K peptides as substrates for *Arabidopsis thaliana* ILK1 (At2G43850) kinase and its D319N mutant in presence of Mn (5 mM).

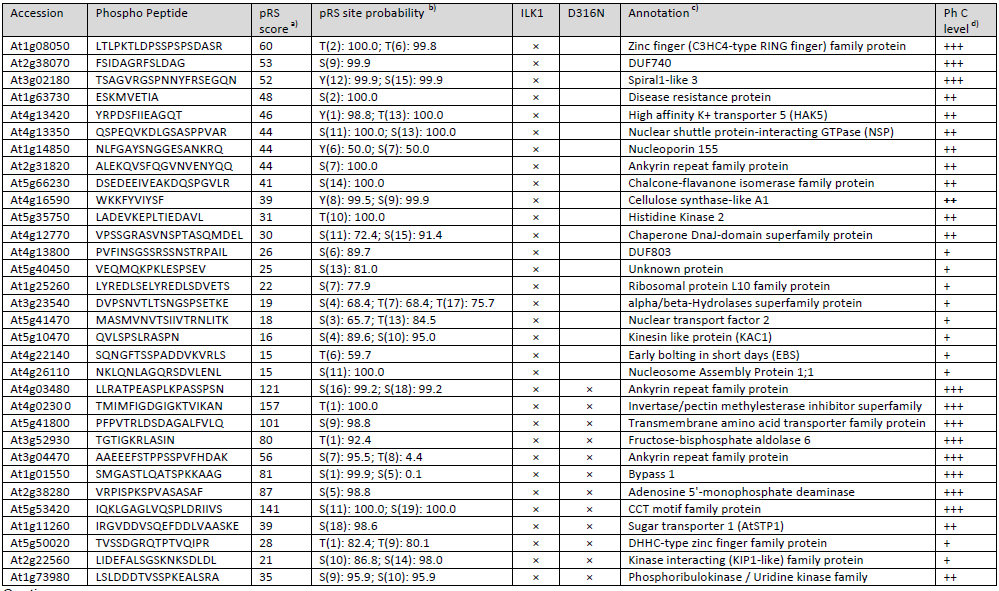

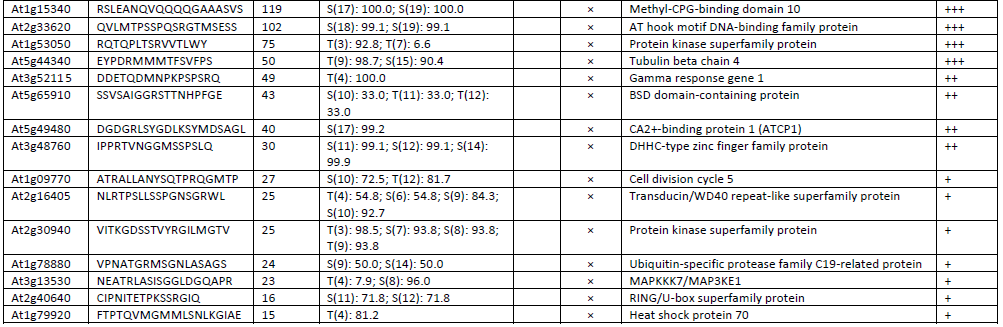

a) pRS score: This peptide score is based on the cumulative binomial probability that the observed match is a random event. The value of the pRS score strongly depends on the data scored, but usually scores above 50 give good evidence for a good PSM.

b) pRS site Probabilities: For each phosphorylation site this is an estimation of the probability (0-100%) for the respective site being truly phosphorylated. pRS Site Probabilities above 75% are good evidence that the respective site is truly phosphorylated.

c) Protein annotation from TAIR database

d) Ph C level: Confidence level of the phosphopeptides

+++ (High): pRS score >50; pRS site probability >75%;

++ (Moderate): pRS score 30-49; pRS site probability 40-74%;

+ (Low): pRS score 15-30; pRS site probability >40%;

**Supplementary Table 4:** Expression of putative ILK1 interactors with significantly altered expression in Arabidopsis roots following flg22 or pep1 treatments. The log2FoldChange reported in Rich-Griffin (2020) are displayed.

| Assay^1^ | Isoform and cofactor^2^ | TAIR ID^3^ | flg22 (epi)^4^ | flg22 (cor) ^4^ | pep1 (epi) ^4^ | pep1 (cor) ^4^ | Annotation^5^ |
| --- | --- | --- | --- | --- | --- | --- | --- |
| KiC | Mn-I | At5g66230 |  |  | -1.60 |  | Chalcone-flavonone isomerase family protein |
| FPM | Mn/Mg-I | AT5G02890 | -1.89 |  | -2.96 | -4.04 | HXXXD-type acyl-transferase family protein |
| FPM | Mn-I | AT2G17680 |  |  |  | -3.01 | DUF241 domain protein |
| KiC | Mn-I | AT5G35750 |  | -1.10 | -2.05 | -2.25 | AHK2 Histidine kinase |
| FPM | Mn/Mg-I | AT2G36410 |  | -1.16 | -1.35 | -2.04 | none |
| FPM | Mn/Mg-I | AT4G25250 |  |  | -2.28 | -1.71 | PMEI4 Pectinesterase inhibitor |
| FPM | Mn/Mg-I | AT3G09260 |  | -0.87 | -1.08 | -1.48 | PYK10 beta-glucosidase |
| KiC | Mn-I | At5g40450 |  | -0.90 |  | -1.46 | none |
| FPM | Mn/Mg-I | AT4G22150 |  |  | -1.09 | -1.37 | Plant UBX domain-containing protein 3 |
| FPM | Mn/Mg-I | AT5G24430 |  |  |  | 0.82 | CDPK-related kinase 4 |
| KiC | Mg-I | AT1G06070 |  |  |  | 1.11 | none |
| KiC | Mg-I,D Mn-D319N | AT1G53050 |  |  |  | 1.38 | Protein kinase superfamily |
| FPM | Mn/Mg-I | AT3G06270 |  |  |  | 1.89 | Protein phosphatase 2C 35 |
| FPM | Mn/Mg-I | AT1G06135 |  |  |  | 2.44 | none |
| KiC | Mn,Mg-I | AT1G08050 | 2.05 | 3.54 | 4.13 | 5.20 | Zinc finger protein |
| FPM | Mn/Mg-I | AT1G30710 |  | 5.59 |  |  | Berberine bridge enzyme-like |
| FPM | Mn/Mg-I | AT1G53625 | 2.67 | 4.07 |  |  | none |
| FPM | Mn/Mg-I | AT5G38310 |  |  | 4.96 |  | none |

^1^Putative ILK1 interactors were identified by KiC (KiC) or functional protein microarray (FPM) in this study.

^2^Two purified ILK1 isoforms were used in the KiC including, ILK1 (I), and ILK1^D319N^ (D). MnCl_2_ (Mn) or MgCl_2_ (Mg) were used in the KiC or both cofactors (Mn/Mg) were used in the functional protein microarray.

^3^The Arabidopsis information resource identification (TAIR) number is indicated.

^4^The log2 fold change reported in Rich-Griffin (2020) is displayed for root epidermis (epi) or cortex (cor+) cell layers comparing MAMP-treated compared to mock-treated roots.

^5^Annotations were derived from the TAIR website (www.arabidopsis.org).

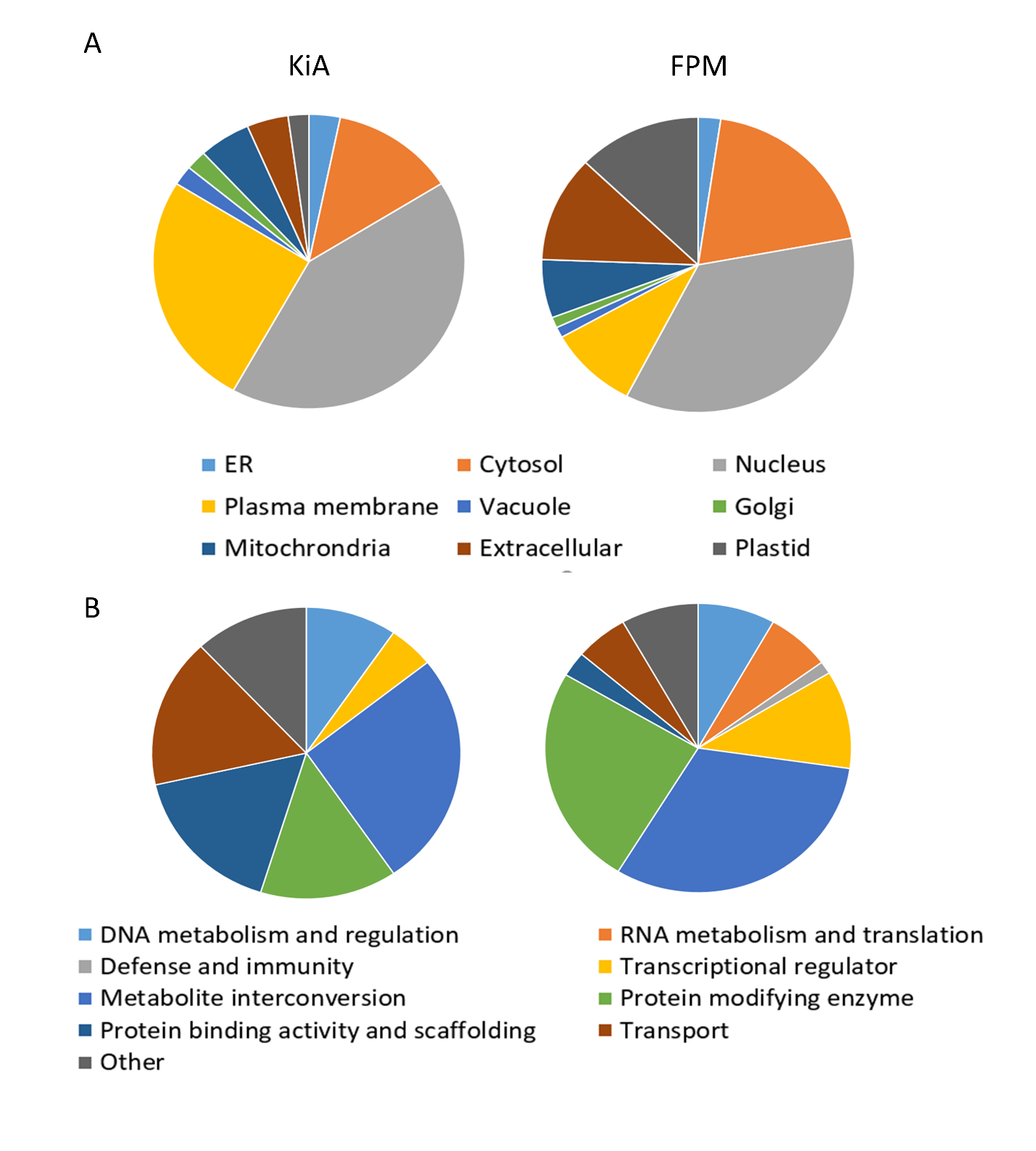

**Supplementary Figure 1:** Localization and protein class of the putative ILK1 targets identified by KiC (KiC) and functional protein microarray (FPM). A) Localization for each protein was identified using the SUBA database (www.suba.live) and can be found in Supplementary Data 1. B) The protein class description was identified using the Panther classification system (pantherdb.org).
